# Alpha-synuclein pre-formed fibrils injected into prefrontal cortex primarily spread to cortical and subcortical structures and lead to isolated behavioral symptoms

**DOI:** 10.1101/2023.01.31.526365

**Authors:** Matthew A. Weber, Gemma Kerr, Ramasamy Thangavel, Mackenzie M. Conlon, Hisham A. Abdelmotilib, Oday Halhouli, Qiang Zhang, Joel C. Geerling, Nandakumar S. Narayanan, Georgina M. Aldridge

## Abstract

Parkinson’s disease dementia (PDD) and dementia with Lewy bodies (DLB) are characterized by diffuse spread of alpha-synuclein (α-syn) throughout the brain. Patients with PDD and DLB have a neuropsychological pattern of deficits that include executive dysfunction, such as abnormalities in planning, timing, working memory, and behavioral flexibility. The prefrontal cortex (PFC) plays a major role in normal executive function and often develops α-syn aggregates in DLB and PDD. To investigate the consequences of α-syn pathology in the cortex, we injected human α-syn pre-formed fibrils into the PFC of wildtype mice. We report that PFC PFFs: 1) induced α-syn aggregation in multiple cortical and subcortical regions with sparse aggregation in midbrain and brainstem nuclei; 2) did not affect interval timing or spatial learning acquisition but did mildly alter behavioral flexibility as measured by intraday reversal learning; 3) increased open field exploration; and 4) did not affect susceptibility to an inflammatory challenge. This model of cortical-dominant pathology aids in our understanding of how local α-syn aggregation might impact some symptoms in PDD and DLB.

## INTRODUCTION

Patients with brainstem alpha-synuclein (α-syn) pathology have motor characteristics of Parkinson’s disease (PD), including rigidity, bradykinesia, and tremor [1]. However, patients with PD dementia (PDD) and dementia with Lewy bodies (DLB), collectively termed Lewy Body dementias (LBD), have diffuse spread of α-syn throughout the brain, including cortical regions [2,3]. These patients often suffer from severe symptoms, including executive and working memory deficits, fluctuations in arousal and attention, hallucinations and psychosis, and increased risk for delirium [3–6]. It is not known whether the spread of α-syn to the cortex causes the symptoms of dementia, such as impaired cognition, or whether cortical α-syn simply reflects a more general process of spread and severity that does not directly affect cortical and cognitive function. To date, there are very few effective therapies approved to manage the debilitating cognitive symptoms of dementia [7,8]; thus, it is critical that we understand the etiological factors of the diverse symptoms seen in LBD.

To investigate the impact of cortical-predominant pathology, we induced α-syn aggregation using human pre-formed fibrils (PFFs) injected in the prefrontal cortex (PFC) of wild-type mice, which can induce aggregation of native α-syn that resembles the pathology described in PDD and DLB [9–12]. Prior literature has extensively studied both mouse and human α-syn injected in the rodent striatum, the predominant model currently used primarily due to its unique ability to induce cell loss in the substantia nigra [11,13]. A few studies have used varying injection locations, including duodenal, pedunculopontine nucleus, nucleus accumbens, olfactory bulb, and others [14–20]. These studies have offered mixed results regarding behavioral deficits and often restrict analyses to 6–7 months post-injection (mpi). Furthermore, only a few studies have investigated PFC PFF injections [21–23]. A prior report from our group found no behavioral effects of PFC PFFs [23]. These animals had a combined viral overexpression/PFF expression strategy, aged animals to 15 months (6 mpi), and examined interval timing, novel object recognition, and open field behavior [23]. In this study, we investigated the effect of PFC PFFs on cortical, subcortical, midbrain, and brainstem α-syn pathology in mice aged up to 2 years (21 mpi) with a broader range of behavioral assays and challenges.

We report four results: 1) PFFs injected in the PFC caused a consistent pattern of moderate-to-severe pathological α-syn aggregation in cortical and subcortical regions with sparse aggregation in midbrain and brainstem nuclei; 2) cognitive behaviors were mildly affected in mice injected with PFFs in the PFC, with no change in interval timing or spatial working memory acquisition, but a small, consistent alteration in behavioral flexibility as measured by intraday reversal learning; 3) mice with cortical PFFs showed greater exploration of the center of an open field, without a change in total distance traveled; 4) mice with cortical PFFs did not have enhanced susceptibility to a delirium-like trigger. Together, these data suggest that aggregation of α-syn in cortical regions impacts specific behaviors even in the absence of midbrain/brainstem inclusion. These results provide insight into cortical synucleinopathy, which is relevant for LBD.

## MATERIALS AND METHODS

### Mice

All experimental procedures were performed in accordance with relevant guidelines by the University of Iowa Institutional Animal Care and Use Committee (IACUC). Wild-type male C57BL/6 mice were received from Jackson Labs (Bar Harbor, ME) at approximately 8–10 weeks of age and acclimated to the animal holding facility for 1–2 weeks. All mice were communally housed on a 12-hour light/dark cycle with *ad lib* access to standard laboratory chow and water, except as noted.

### Monomer and fibril preparation

Monomeric alpha-synuclein and human pre-formed fibrils (PFFs) were a contribution from Dr. Andrew West and prepared according to previously used protocols [23,24]. On the day of surgery, aliquoted monomers and PFFs were thawed and kept on ice immediately prior to surgical procedures. Potentially contaminated surfaces were then thoroughly cleaned with 10% bleach.

### Surgical Procedures

Mice were anesthetized under 4.0% isoflurane and surgical levels of isoflurane (1.5–3.0%) were sustained for the duration of the surgery (SomnoSuite, Kent Scientific, Torrington, CT, USA). Analgesic (carprofen) and local anesthetic (bupivacaine) were administered prior to the start of surgery. An incision was made along midline and bilateral burr holes drilled above the prefrontal cortex (PFC), and a Hamilton syringe was lowered to the target coordinates (AP +1.8, ML ±0.5, DV -1.5). Mice were assigned to bilateral microinjections of either 2 µL monomers or 2 µL PFFs. Monomeric controls or PFFs were infused over 20 minutes (0.1 µl/min; Legato 130 Syringe Pump, kd Scientific, Holliston, MA, USA), with a 5-minute wait period before removing the needle. The incision was then sutured, and mice were moved to a clean cage with *ad lib* access to food and water. Behavioral testing began approximately 12 months later (Fig 1).

**Figure 1.**
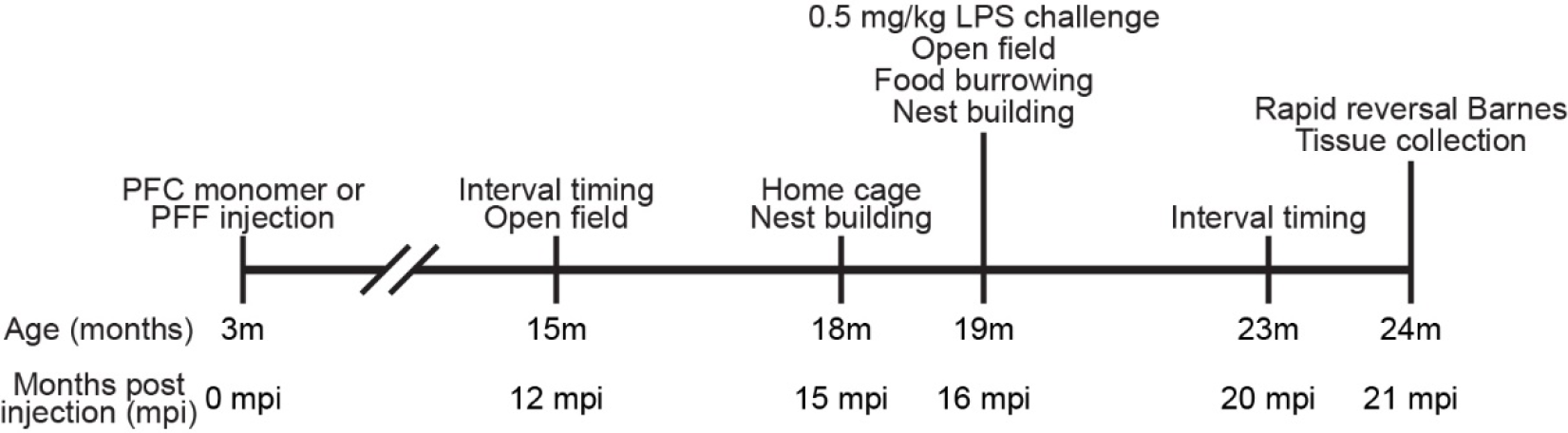
Experimental timeline.

### Interval Timing Switch Task

All mice were trained to perform an interval timing switch task described in detail previously [25–28]. Interval timing behavior was assessed on two occasions: 1) 12 months post-injection (mpi); 2) 20 mpi. Briefly, food-restricted mice are trained in standard operant chambers (MedAssociates, St. Albans, VT) equipped with two nosepoke response ports (left and right) flanking a reward hopper and another nosepoke response port on the back wall. Either the left or right nosepoke was designated for short trials and the contralateral nosepoke for long trials (counterbalanced across mice and experimental groups). Trials are initiated by a response at the back nosepoke, which generated identical cues for both the left and right nosepokes. In short trials, mice receive a reward for the first response after 6 seconds. In long trials, mice receive a reward for the first response after 18 seconds. Because cues are identical for both short and long trials, the optimal strategy is to start responding at the short nosepoke and switch to the other nosepoke to receive a reward if enough time passed without reward.

### Rapid reversal Barnes maze

A modified Barnes maze was used to evaluate spatial learning acquisition and behavioral flexibility using a modified Barnes maze adapted from established protocols [29,30], with the following notable exceptions: 1) the spatial target was changed every day to a new random location and every 4 trials to an opposite location in order to assess rapid acquisition of a spatial target (first 4 trials) and intraday reversal learning (last 4 trials); 2) to minimize the use of olfactory cues, every hole was equipped with its own 3D printed escape box “shuttle” such that no shuttle was ever re-used during a session, and the maze itself was also rotated. A plastic grate made of identical material prevented entry to incorrect shuttles; 3) following a correct response (entry), the shuttle was connected by tunnel with the home cage as a potent reward, during a 3-minute inter-trial interval; 4) a 40-hole circular Barnes maze apparatus was used in which only 10 holes were available. Total distance traveled was captured with overhead cameras and analyzed using ANY-Maze 7.14d (Stoelting, Wood Dale, IL).

### Open field

Open field activity was used to assess exploration and activity levels on two occasions (12- and 16 mpi), using methods described previously [31,32]. On each occasion, mice were habituated to behavior rooms for at least 30 minutes prior to testing. Mice were placed in the center of an empty open field arena (40 cm × 40 cm × 30 cm) and allowed to explore uninterrupted for 10 minutes. Mice were returned to their home cage immediately, and the open field arenas were thoroughly cleaned with 70% ethanol and allowed to air dry between trials. Rescue disinfectant (0.5% hydrogen peroxide) was used between cohorts. Total distance traveled and thigmotaxis (distance traveled outside a 25.5 × 25.5 cm central square divided by total distance traveled) were analyzed using ANY-Maze 7.14d.

### Home cage activity

Home cage activity was assessed in parallel with nest building and food burying (described below) at 15 mpi. Overhead cameras captured morning and nighttime activity using an infrared camera. ANY-Maze 7.14d software was then used to quantify periods of activity and rest over a one-hour period during the morning (∼6–7 am) and night (∼2–3 am). Midday activity could not be quantified due to unexpected interference with lighting through cage bars. Periods of immobility were defined as periods 40-seconds or longer during which ANY-Maze did not detect movement. Percent immobile was then defined as the number of seconds the mouse spent immobile compared with total seconds tracked. Bout length was based on prior published data using EEG suggesting that periods of immobility greater than 40-seconds are likely to be sleep bouts [33].

### Nesting

Nest building was used to approximate general health and welfare, using slightly modified methods described previously [34]. Nesting behavior for each mouse was assessed on two occasions (15- and 16 mpi). At 15 mpi, three different materials (shredded paper, paper twist, and pressed cotton square) were provided on separate days. At 16 mpi, only the paper nest building material was provided. Photographs were taken of each nest and scored by blinded observers on a scale of 1–5 according to previous literature [34,35].

### Food burying

Food burying was assessed in parallel with nest building at 16 mpi. Pre-weighed food was provided in a glass jar equipped with a PLA 3D printed cone-shaped hopper to prevent mice from nesting in the jar. The jar was weighed the following day to calculate food buried or eaten.

### Lipopolysaccharide injection challenge

Systemic infection increases risk of developing delirium and dementia in humans [36–38] and peripheral injections of lipopolysaccharide (LPS) are widely used as a model of systemic infection and resultant behavioral changes in rodents [39–41]. At 16 mpi, 0.5 mg/kg LPS (Sigma #L2880, St. Louis, MO) diluted in 0.9% sterile saline was injected intraperitoneally (IP) in five mice with PFC monomers and five mice with PFC PFFs. This dose of LPS was chosen as previous studies demonstrated accelerated decline in rodent models of tauopathy and prion disease [42,43]. The remaining three mice with PFC monomers and three mice with PFC PFFs were injected with equivalent volumes of saline. Behavior in the open field, food burying assay, and nest building was then assessed. At 21 mpi, a lower dose (0.25 mg/kg LPS) was injected IP to all mice to examine acute effects of the challenge in the open field and rapid reversal Barnes maze 48 hours prior to perfusion (Fig. S2).

### Histology

Mice were anesthetized with ketamine (100 mg/kg IP) and xylazine (10 mg/kg IP) and transcardially perfused with phosphate-buffered saline (PBS) and 4% paraformaldehyde (PFA). Brains were fixed in 5% PFA overnight and immersed in 30% sucrose for approximately 48 hours. Brains were then serially sliced into 40-μm coronal sections using a cryostat (Leica Biosystems, Deer Park, IL). Brain sections were blocked for one hour in 2% normal goat serum (NGS) in PBST (0.3% Triton X-100 in 1x PBS) and then incubated for approximately 20 hours at 4°C in either rabbit anti-α-synuclein (α-syn) phosphorylated serine 129 (P-S129; Abcam #EP1536Y, Cambridge, United Kingdom) diluted to 1:1000 in blocking solution or rabbit anti-tyrosine hydroxylase (TH; Millipore #AB152, Burlington, MA) diluted to 1:1000 in blocking solution. Sections were then washed in PBS three times and incubated overnight at 4°C in goat anti-rabbit IgG (H+L) Alexa Fluor 647 secondary antibody (Invitrogen #A11036, Waltham, MA) diluted to 1:1000 in blocking. Sections were washed three more times. To stain for nissl, sections were washed in PBST for 10 minutes and two additional 5-minute washes in PBS. Sections were incubated in NeuroTrace™ 435/455 (Thermo Fisher Scientific Inc. #N21479, Waltham, MA) diluted 1:300 in PBS for 20 minutes at room temperature and then washed in PBST for 10 minutes and twice in PBS for 5 minutes each. After one more wash at room temperature for 2 hours, sections were mounted with ProLong Diamond Antifade Mountant (Invitrogen #P36961) on Superfrost microscope slides (Fisher Scientific).

All sections were imaged using VS-ASW-S6 imaging software (Olympus, Center Valley, PA) and an Olympus Slide Scanner (Olympus VS120) with a 20x objective. To characterize pathological spread of α-syn, we semi-quantitively graded the degree of P-S129 inclusions in multiple regions of interest (ROI; see Fig. 2 for complete list) referenced from Franklin & Paxinos, 2008. The α-syn pathology was graded using the rubric images from previously published literature [45]. Briefly, pathology was assigned on a scale of 0–4 separately for neurites and somas, with the following general guidelines, in addition to published rubric, to assist and calibrate blinded grading; 0: no soma or neurite P-S129 positive staining; 1: sparse/light soma or neurite staining (1–3 somas or neurites per 750 µm square area within ROI); 2: mild burden (4 or more somas or neurites per 750 µm, with definitive areas showing no P-S129 staining); 3: dense burden (>10 soma or neurites per 250 µm); 4: heavy burden (>20 somas or neurites per 250 µm). For every brain, one out of every six 40 µm sections were graded for α-syn inclusion. Results from brain regions that spanned multiple sections were averaged across sections and hemispheres, even if some sections contained no pathology.

**Figure 2.**
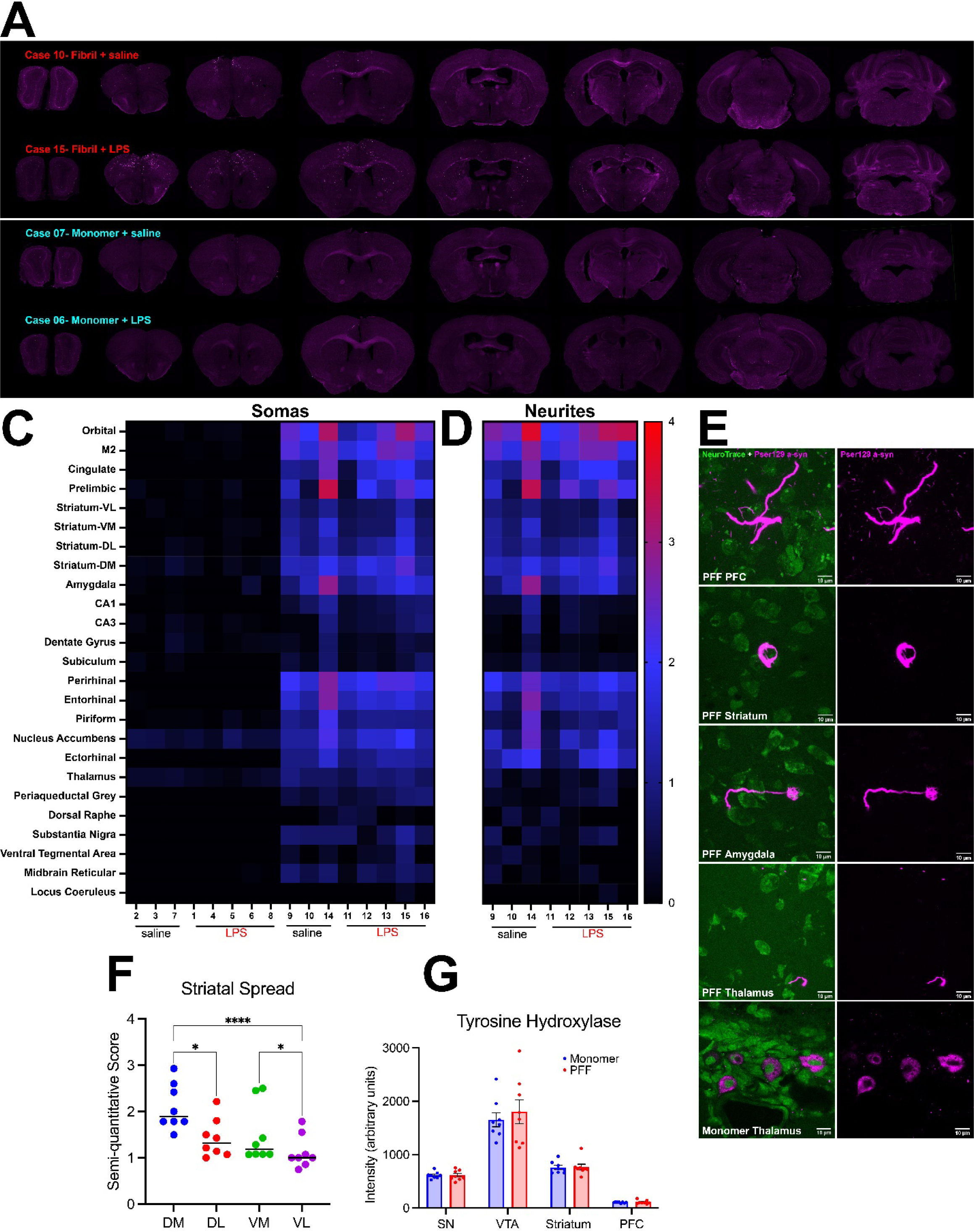
Evaluation of pathology in monomer control and pre-formed fibril (PFF) injected mice. A) Top: representative high-resolution images across the entire rostral-caudal aspect of the brain in two mice treated with PFFs (one saline and one LPS treated at 16 mpi, sacrificed at 21 mpi). Bottom: representative images of two monomer control mice (one saline and one LPS treated at 16 mpi, sacrificed at 21 mpi). Semi-quantitative scores of phospho α-syn (p-α-syn) positive **C)** somas and **D)** neurites of 25 brain regions, with each column representing an individual mouse, averaged over the entire rostral-caudal extent for each brain region examined. Neurites for monomer mice are not displayed as there were zero p-α-syn positive neurites in any region of any mouse. **E)** Confocal images showing examples of fibrillar pathology in multiple brain regions from PFF injected mice (panels 1–4). Monomer mice showed rare p-α-syn positive puncta, without clear fibrillar morphology (panel 5). No fibrillar or neurite pathology was found in monomer mice using confocal microscopy. **F)** Averaged semi-quantitative score of pathological α-syn within subregions of the striatum restricted to 0–1.7 mm bregma. Horizontal bars indicate group medians. * p < 0.05, **** p < 0.0001 **G)** Tyrosine hydroxylase intensity (arbitrary units) from 4 different brain regions PFF treated mice: substantia nigra (SN), ventral tegmental area (VTA), striatum, and prefrontal cortex (PFC). Tyrosine hydroxylase intensity expressed as mean ± SEM, and each dot represents a single mouse.

### Statistics

All data was analyzed using custom written MATLAB routines and Graphpad Prism.

All tests with ordinal outcomes were analyzed using appropriate non-parametric statistics. Pathology scores (ordinal) were analyzed using Friedman Rank Sum test. Tyrosine hydroxylase values (quantitative intensity) were analyzed using a repeated-measures ANOVA. Behavioral tasks were analyzed using separate unpaired t-tests or repeated measures ANOVA. Separate nest building scores (ordinal) were analyzed using separate Mann-Whitney *U* tests. Behavioral tasks at 16 mpi – 0.5 mg/kg LPS challenge – were analyzed using two- or three-way ANOVA or Friedman Rank Sum test (nest building only) followed by separate Mann-Whitney *U* tests corrected for multiple comparisons.

## RESULTS

### Pathology

Injections of PFFs in the PFC led to α-syn aggregation in the injected cortical area as well as several downstream regions. We found clear and dense pathological spread of α-syn in orbitofrontal, secondary motor M2, perirhinal, entorhinal, piriform, nucleus accumbens, amygdala, and thalamus (Fig. 2C-D). Subregions of the striatum were differentially affected by PFFs injected in the PFC (Fig. 2F). A Friedman test was conducted and there was a statistically significant difference in α-syn aggregation between subregions of the striatum (Χ^2^(3) = 22.2, p < 0.0001). Post-hoc analyses using Dunn’s multiple comparisons test revealed significant differences between dorsomedial and dorsolateral striatum (*p* = 0.04), dorsomedial and ventrolateral striatum (*p* < 0.0001), and ventromedial and ventrolateral (*p* = 0.04). Unlike other subcortical regions, subregions of the hippocampus were considerably less dense, and similar sparse-to-absent pathological α-syn aggregation was observed in regions of the midbrain and brainstem.

We also quantified tyrosine hydroxylase positive (TH+) fluorescence levels in the PFC, striatum, ventral tegmental area, and substantia nigra (Fig. 2G), similar to prior methods established in the lab [28]. As expected, there was a statistically significant difference in TH+ fluorescence levels between regions (Fig. 3E; two-way repeated-measures ANOVA; main effect of region (F_(1.105, 15.47)_ = 122.9; *p* < 0.0001), but we found no significant change in TH+ fluorescent levels in PFF-injected mice compared with controls in any of the four regions (main effect of treatment (F_(1, 14)_ = 0.2078; *p* = 0.66). Further, we did not find any differences in markers for astrocyte (GFAP) or microglia (IBA-1) activation (Fig. S2D).

**Figure 3.**
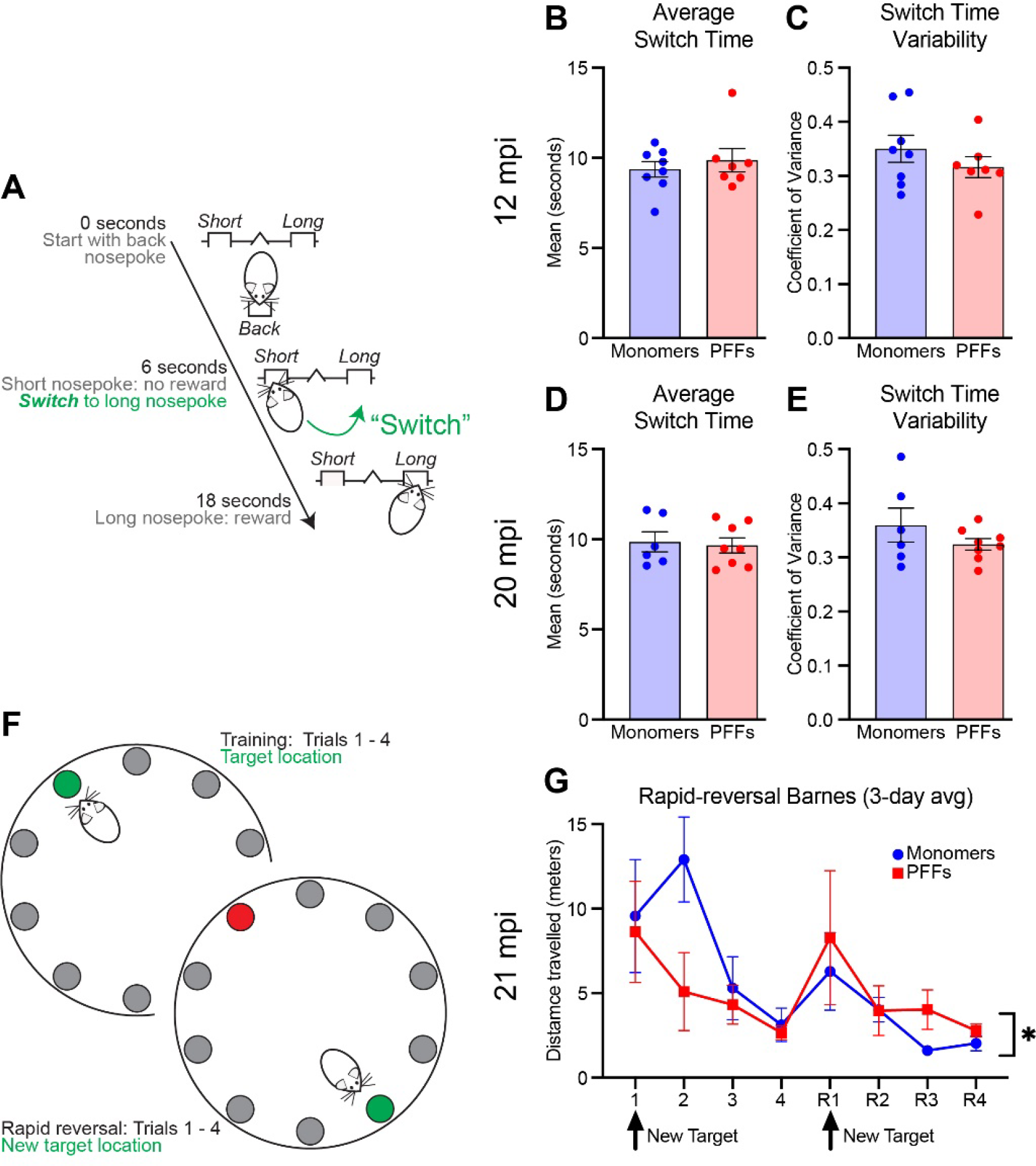
Interval timing and reversal learning in monomer control and PFC PFF mice. A) Diagram for the interval timing switch task outlining optimal performance during long trials. **B)** Mean switch time and **C)** switch time coefficient of variance during the interval timing task at 12 mpi. Data from 8 monomer (blue) and 7 PFF (red) mice. **D)** Mean switch time and **E)** switch time coefficient of variance during the interval timing task. Data from 6 monomer (blue) and 8 PFF (red) mice. Each dot represents a single mouse. **F)** Diagram for the rapid reversal Barnes maze. **G)** Average total distance travelled averaged across three-days. There was a main effect of trial and trial × treatment interaction. * p < 0.05, but post-hoc testing at specific trials did not survive multiple comparisons. Data from 6 monomer (blue) and 6 PFF (red) mice. Data are expressed as mean ± SEM.

### Cognitive behaviors: Interval timing and behavioral flexibility

Interval timing requires mice to estimate an interval of several seconds and requires working memory for temporal rules as well as attention to the passage of time. Our prior work showed no deficits in interval timing using a fixed-interval timing task at 15 months of age (6 months post injection (mpi) of PFFs) [23]. Here, we sought to investigate interval timing performance in mice at 12+ mpi, using an interval timing switch task that enables us to track mean temporal estimates as well as the variability of temporal estimates. We found that at 12 mpi, mice injected with PFFs showed no difference in mean switch times (shifting early or late as an estimate of timing accuracy; Fig. 3A; unpaired t-test; *t*(13) = 0.66; *p* = 0.52) and no difference in switch time coefficient of variability (estimate of timing precision; Fig. 3B; unpaired t-test; *t*(13) = 1.05; *p* = 0.31). One PFF injected mouse was excluded because no trials initiated or rewards obtained, likely due to a lack of motivation to perform the task. We repeated interval timing switch task training at 20 mpi. Two monomer-treated mice were sacrificed at 19 mpi due to age-related health concerns per the University of Iowa Office of Animal Resources guidelines, so this round of interval timing training included six monomer and eight PFF mice. Similar to interval timing at 12 mpi, we found no difference in mean switch times (Fig. 3C; unpaired t-test; *t*(12) = 0.29; *p* = 0.78) or the coefficient of variability of switch times (Fig. 3D; unpaired t-test; *t*(12) = 1.20; *p* = 0.25). We found no difference in the proportion of rewarded trials at either timepoint (Fig. S1A & B). Together, these results suggest that PFC PFFs do not affect interval timing behavior even at 20 mpi.

After interval timing training, two mice treated with PFFs were sacrificed due to age-related health concerns. Therefore, six monomer and six PFF mice remained. To test learning and behavioral flexibility, all remaining mice were tested using a rapid reversal Barnes maze at 21 mpi. PFF mice showed an altered pattern of behavior flexibility, with PFF mice learning quickly during the initial trials, but then travelling a greater distance to reach the target compared with control mice during the reversal phase (Fig. 3E; two-way repeated-measures ANOVA; main effect of trial (F_(2.145, 19.31)_ = 6.71; *p* = 0.005) and trial × treatment interaction (F_(7, 63)_ = 2.17; *p =* 0.049); post-hoc analyses revealed no significant differences when corrected for multiple-comparisons). A similar pattern of behavior during the reversal phase, although not statistically different, was observed during a single day rapid reversal Barnes protocol at 17 mpi (Fig. S1C).

### Open field

We assessed exploration and activity levels in the open field. At 12 mpi, mice injected with PFFs in the PFC did not differ from monomer controls in distance traveled (meters) over the 10-minute test (Fig. 4A; unpaired t-tests; *t*(13) = 0.35; *p* = 0.73), but did display reduced thigmotaxis, showing more exploration of the center of the arena (Fig. 4B; unpaired t-tests; *t*(13) = 2.49; *p* = 0.03). These results may suggest greater exploration or reduced anxiety-like behavior.

**Figure 4.**
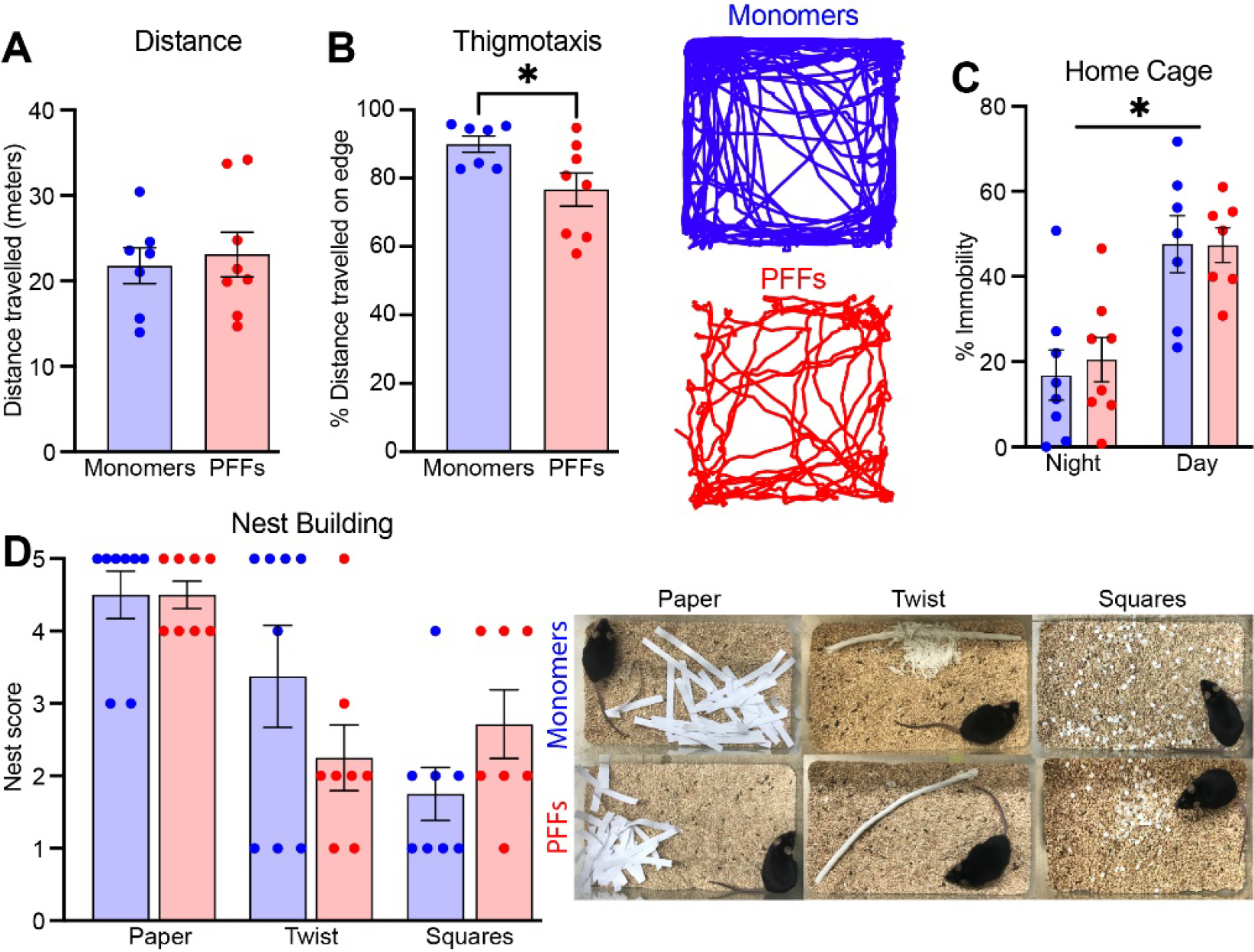
Open field, home cage activity, and nest building in monomer control and PFC PFF mice at 12–15 mpi. A) Distance travelled and **B)** thigmotaxis during the open field test. The right panel is an example trace from one monomer and one PFF mouse during the open field assay. **C)** Percent immobility time between ∼2–3 am (night) and ∼6–7 am (morning). **D)** Nest building score for three different materials (shredded paper, cotton twist, and precut cotton squares) with representative images from each material shown in the right panel. Data from 8 monomer (blue) and 7-8 PFF (red) mice. All data are expressed as mean ± SEM, and each dot represents a single mouse. * p < 0.05

### Home cage activity and nest building (15 mpi)

To assess general health and welfare as well as naturalistic behaviors, we used assays to examine home cage activity and nest building. At 15 mpi, mice injected with PFFs in the PFC did not differ on amount of home cage activity during either the light or dark cycle (Fig. 4C; mixed-effects two-way ANOVA; main effect of treatment F_(1, 14)_ = 0.04; *p* = 0.84), but there was a significant main effect of time of day (F_(1, 12)_ = 50.45; *p* < 0.0001). At the same time-point, there were no differences in nest building score across three different types of material (Fig. 4D; Mann-Whitney *U*; paper *p* = 0.61; twist *p* = 0.42; squares *p* = 0.18). Altogether, these data suggest that PFC PFFs do not reduce naturalistic innate behaviors or overall well-being.

### Peripheral immune challenge (16 mpi)

We tested if mice previously injected with PFF were more susceptible to peripheral immune activation by lipopolysaccharide (LPS) injections. At 16 mpi, we injected five mice with PFC PFFs and five mice with PFC monomers with 0.5 mg/kg LPS. The remaining three mice in each group received equivalent volumes of saline. We then tested behavior in the open field, food burying, and nest building assays. Although LPS caused a significant decrease in distance travelled (Fig. 5A; two-way ANOVA; F_(1, 12)_ = 44.82; *p* = < 0.0001) and food burying (Fig. 5C; three-way ANOVA; F_(1, 12)_ = 25.64; *p* = 0.0003), we found no evidence of an enhanced response in PFF injected mice (Fig. 5A; distance travelled F_(1, 12)_ = 1.33; *p* = 0.27; Fig. 5B; two-way ANOVA; thigmotaxis F_(1, 11)_ = 0.52; *p* = 0.49; Fig. 5C; food burying F_(1,12)_ = 1.32; *p* = 0.27). Unlike 12 mpi, there was no significant effect of PFFs on thigmotaxis (F_(1, 11)_ = 2.46; *p* = 0.15), possibly due to a lack of statistical power, as the pattern remained similar even in the presence of LPS. Similarly, although LPS impaired nest building (Χ^2^(4) = 12.18, p = 0.02), this response was not enhanced in PFC PFF mice (Fig. 5D; 1-day post LPS injection; Mann-Whitney *U*; *p* = 0.16). These results suggest that PFC PFF mice are not more susceptible to peripheral immune challenge. This is further supported by a final lower dose challenge of 0.25 mg/kg LPS given at 21 mpi. PFC PFF mice were not more susceptible to LPS challenge in the open field or rapid reversal Barnes maze (Fig. S2).

**Figure 5.**
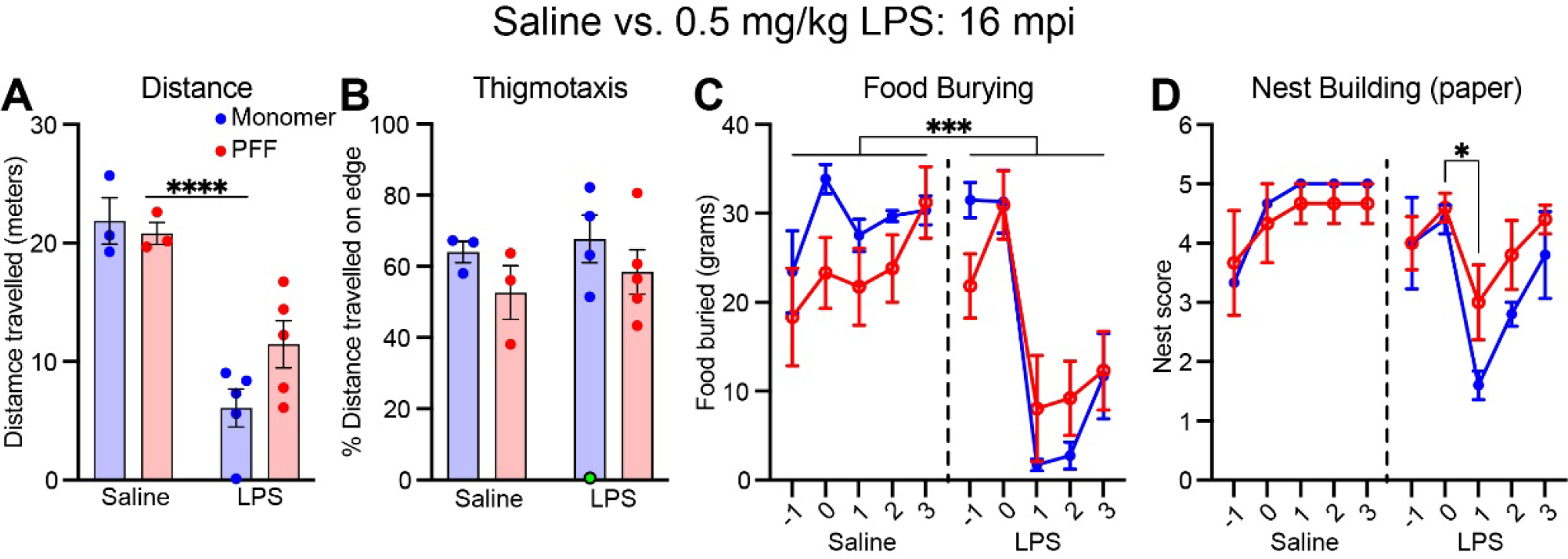
Open field, food burying, and nest building during 0.5 mg/kg LPS challenge. A) Distance travelled and **B)** thigmotaxis during the open field after either saline or lipopolysaccharide (LPS) injection. There was a main effect of LPS on distance traveled, but this effect was not different for monomer and PFF mice. One monomer mouse treated with LPS was excluded from thigmotaxis analysis as no movement was recorded during the assay. **C)** Amount of food buried (in grams) during the food burying assay and **D)** nest building score (shredded paper only) for 2 days before and 3 days after either saline or LPS injection. All data are expressed as mean ± SEM, and each dot represents a single mouse. * p < 0.05; *** p < 0.001; **** p < 0.0001

## DISCUSSION

We tested the effects of prefrontal cortex (PFC) human pre-formed fibrils (PFFs) in aged mice on multiple behavioral assays and then evaluated spread of α-synuclein (α-syn) aggregation in cortical, subcortical, and brainstem regions. First, we found that PFC PFF injection triggered widespread cortical and subcortical α-syn aggregation, without significant spread to brainstem regions, allowing evaluation of the behavioral impact of α-syn limited to these regions. Second, and in line with our prior work, we found no reliable deficits in interval timing, but we did find mild alterations in reversal learning. Third, we also found changes in exploration without changes in distance travelled in the open field assay. Finally, we found no effects of inflammatory challenge. Taken together, these data contribute to understanding potential consequences of cortical α-syn, which is important for efforts investigating the role of this pathological finding in synucleinopathies such as Parkinson’s disease dementia (PDD) and dementia with Lewy bodies (DLB).

The rodent PFC is a highly connected brain region, sending projections to multiple cortical, subcortical, midbrain, and brainstem nuclei [46,47]. Our prior work suggested that human PFFs injected into PFC cause a consistent but spatially limited pattern of α-syn pathology including PFC, temporal cortex, and striatum. The consequences of α-syn outside of the brainstem, as occurs in PDD and DLB, are still relatively unknown. Using the model of cortically-injected PFFs helps tease out the consequences of cortical/subcortical α-syn pathology in the absence of dramatic loss of neurons in brainstem regions.

Using semi-quantitative analysis of 25 brain regions, we demonstrate dense pathological spread of α-syn in multiple cortical regions such as the orbitofrontal, secondary motor M2, perirhinal, entorhinal, and piriform. A consistent pattern of α-syn was also observed in subcortical regions, including the striatum, nucleus accumbens, amygdala, and thalamus. We note that there is differential aggregation of α-syn in subregions of the striatum, matching the connectivity of our prefrontal injection site and in line with prior work from Gabbott and colleagues (2005). We targeted the dorsal prelimbic and anterior cingulate subregions of the PFC, which contain a greater number of neurons that project to the dorsal striatum compared to the ventral prelimbic and infralimbic subregions of the PFC that contain more ventral striatal projecting neurons [48]. We also note that, unlike other subcortical regions, the subregions of the hippocampus were relatively spared from α-syn aggregation. This finding was again in line with prior literature suggesting that cortical input to the hippocampus passes through the perirhinal and entorhinal cortices first [49], two areas that show dense α-syn deposition here. While there was clear α-syn in cortical and subcortical regions, we found little or no α-syn pathology in midbrain and brainstem nuclei, including substantia nigra, locus coeruleus, and dorsal raphe. Together, these findings extend our prior work to show that the aggregation induced in cortical and subcortical regions by human PFFs injected into PFC are not cleared over the course of aging, as has been reported in some brain regions [50].

Notably, prior work from our group has shown that human PFFs injected in the PFC does not affect interval timing or open field behavior 6 mpi in mice aged 15 months [23]. By contrast, prior work has shown that people with PD display reliable deficits in interval timing [51–54] and that the PFC is crucial for interval timing behavior [51,55,56]. The data presented here, and our prior report demonstrate that PFC PFFs are not sufficient to affect interval timing. There are several interesting hypotheses that could explain these results; 1) midbrain dopaminergic cell loss, which is absent in this model, may be the main driver for interval timing deficits seen in PD and mouse models [28,51]; 2) human PFFs may predominate in a PFC cell type that is unnecessary for interval timing; 3) cortical cells may function even in the setting of α-syn aggregation, either through cellular or circuit-level compensation.

While α-syn aggregates were widespread and dense in cortical and subcortical regions, mice that received PFC PFF injections displayed only mild behavioral deficits. PFF mice did not display any difference in measures of interval timing, including variability of or mean switch times. This result in is line with prior work from our group [23] and suggests that PFC PFFs and the resulting α-syn aggregation is not sufficient to induce an interval timing deficit, even up to 20 mpi. We also found that PFF mice displayed reduced thigmotaxis at 12 mpi, with the same trend observed at 16 mpi, which may suggest a difference in explorative and anxiolytic behaviors in the open field as a result of PFC PFF injections.

Further, we also found that PFF mice showed rapid spatial acquisition but required a longer search route to locate the target exit hole during the reversal phase of a daily spatial training Barnes maze task at 21 mpi, suggesting altered behavioral flexibility.

This study has several limitations. First, our control animals received monomer injections, which has been shown previously to occasionally lead to mild and transient aggregation [57]. We did note some mild p-α-syn positive puncta, but morphology was distinct and punctate, and may have been caused by aging alone. Fibrillar and “lewy-neurite-like” structures were only seen in PFF injected mice, and this was confirmed by confocal microscopy. Second, we did not investigate cell-type preference for human PFF-induced α-syn pathology which could impact the response of the PFC and may be different from the response to alternate models, such as brain homogenate-derived α-syn fibrils; for example, recent publications have suggested human PFFs also induce significant glial inclusions [58,59]. Third, we did not include a group of young mice to determine whether our lack of behavioral change was primarily due to age-related decline in cognitive function or interval timing, which has been observed previously [60,61], although the interval timing behavior presented here is similar to previously published work from our group using healthy, young C57BL/6 mice [26]. Fourth, our measures of interval timing and behavioral flexibility do not fully interrogate cognitive dysfunctions related to PDD and LBD and it is possible that other cognitive behaviors such as attention or impulse control were negatively affected by PFC PFF injections. Fifth, behavioral experiments at the later timepoints are underpowered to detect subtle differences due to age-related loss. Despite these limitations, due to careful pathological confirmation of each mouse in this two-year study, we believe these results inform future research and are important for understanding this model and considerations for testing other models to induce aggregation. Finally, this study was limited to male mice. Recent evidence suggest female mice may show reduced pathology compared with male mice [62]. Human males are more likely to be diagnosed with PD and DLB [63–65]. Understanding potential differences in pathology due to sex is an essential goal of future research and fully powered studies of both sexes are needed.

In summary, our results describe widespread cortical and subcortical α-syn aggregation from PFC PFFs with mild behavioral alterations that were not modulated by an inflammatory challenge. The lack of brainstem involvement is notable and could provide insight into the brain networks that drive behavioral deficits in PD, PDD, and DLB.

## ACKNOWLEDGMENTS

This work was supported by NIH R01s MH116043, NS120987 to NSN and NIH K08 NS109287 and the Iowa Neuroscience Institute to GMA. This work was also supported by NIH P20NS123151 to NSN and GMA. We thank Dr. Andrew West for generously providing the pre-formed fibrils and Travis Larson for assisting with behavioral experiments.

## DATA AVAILABILITY

All raw data are available at https://aldridge.lab.uiowa.edu/open-science.

## CODE AVAILABILITY

All code is available at https://aldridge.lab.uiowa.edu/open-science.

## AUTHOR CONTRIBUTIONS

MAW, HAA, QZ, NSN, and GMA designed the experiments. MAW, MMC, HAA, GK, OH, and GMA performed all experiments. GK, RT, JG, and GMA performed histological analyses. MAW and GMA performed all statistical analyses. MAW and GMA wrote the manuscript, and all authors reviewed and revised the manuscript.

## COMPETING INTERESTS

The authors have no conflict of interest to report.

## SUPPLEMENTAL MATERIALS

**Figure S1.**
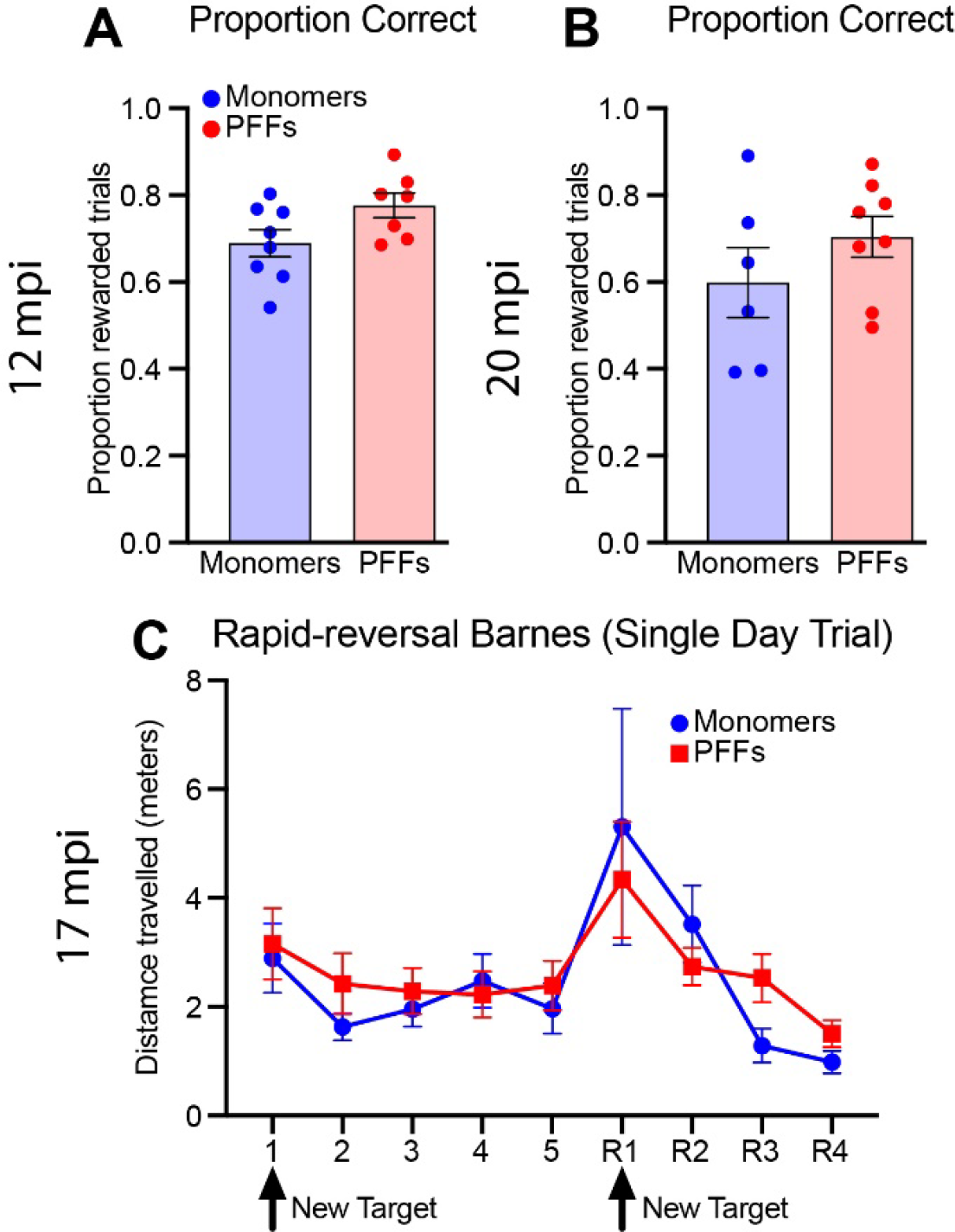
Additional measures of interval timing performance and behavioral flexibility. Proportion of rewarded trials during the interval timing switch task at **A)** 12 mpi and **B)** 20 mpi. Unpaired t-tests revealed no differences in the proportion correct at either timepoint (12 mpi: *t*(13) = 2.03; *p* = 0.06; 20 mpi: *t*(12) = 1.2; *p* = 0.25) **C)** Average total distance travelled during a single day rapid reversal Barnes maze protocol. Compared to the multiple day Barnes maze protocol at 21 mpi, PFC PFF mice show a similar, but non-significant, altered pattern of behavioral flexibility with a greater distance travelled to reach the target during the reversal phase (two-way repeated-measures ANOVA; trial: F_(2.182, 30.55)_ = 4.232; *p* = 0.02; treatment: F_(1, 14)_ = 0.2317; *p* = 0.64) . All data are expressed as mean ± SEM, and each dot represents a single mouse.

**Figure S2.**
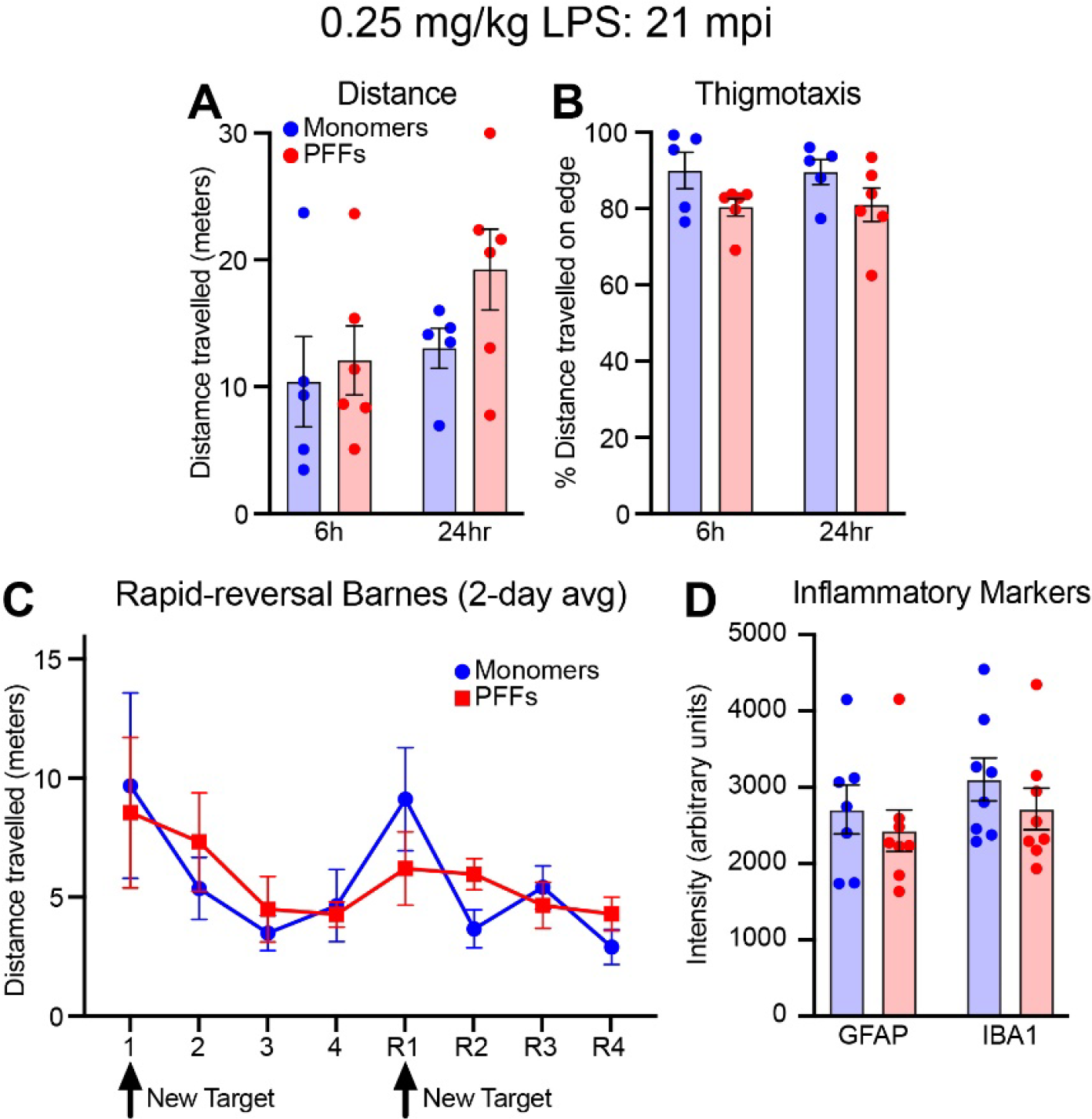
Lower dose lipopolysaccharide (LPS) challenge (0.25 mg/kg) at 21 mpi. A) Distance travelled and **B)** thigmotaxis during the open field test. PFC PFF injected mice did not differ from controls in distance travelled (meters) either 6- or 24-hours post LPS injection (two-way repeated-measures ANOVA; timepoint: F_(1, 9)_ = 5.115; *p* = 0.05; treatment: F_(1, 9)_ = 1.272; *p* = 0.29). However, PFC PFF mice did display a non-significant trend for reduced thigmotaxis (two-way repeated-measures ANOVA; timepoint: F_(1, 10)_ = 0.003; *p* = 0.96; treatment: F_(1, 9)_ = 4.096; *p* = 0.07), regardless of timepoint post injection (F_(1, 10)_ = 0.003; *p* = 0.96). **C)** Average total distance travelled averaged across two-days post LPS of the rapid reversal Barnes maze (two-way repeated-measures ANOVA; trial: F_(2.626, 21.01)_ = 2.667; *p* = 0.08; treatment: F_(1, 8)_ = 0.034; *p* = 0.86). **D)** Fluorescent intensity (arbitrary units) of GFAP and IBA1 in whole striatum. All data are expressed as mean ± SEM, and each dot represents a single mouse.

## Notes

**Funding:** This work was supported by NIH R01s MH116043, NS120987 to NSN and NIH K08 NS109287 and the Iowa Neuroscience Institute to GMA. This work was also supported by NIH P20NS123151 to NSN and GMA.

**Conflict of Interest:** There are no conflicts of interest.

### Competing Interest Statement

The authors have declared no competing interest.

### Summary of Updates

Minor revision to title and author name. Statistics clarified. Figure 2 and 5 updated. Supplemental figures added.

